# Systematic comparison of colocalization methods using protein quantitative trait loci

**DOI:** 10.1101/2025.11.07.686776

**Authors:** Seongwon Hwang, Jeffrey Pullin, Chris Wallace, John Whittaker, Stephen Burgess

## Abstract

Colocalization is frequently performed as a step to triage findings from genetic investigations linking molecular and disease data. However, the reliability and consistency of the various colocalization methods is not well understood. In particular, it is unclear whether a non-colocalization result should be taken as a definitive sign that two traits do not share a genetic cause in the region of interest, or merely as suggestive evidence. We use protein QTL data to benchmark four colocalization methods (coloc, coloc-SuSiE, prop-coloc, and colocPropTest), considering whether the methods conclude there is colocalization between datasets representing associations with the same protein. We consider a baseline scenario in which the associations come from the same dataset split in half at random, and scenarios in which the associations are with the same protein, but measured on different platforms or in different populations. In the baseline scenario, all methods report colocalization for the majority of proteins. In other scenarios, methods do not consistently report colocalization, they often report non-colocalization, and they often disagree. In the worst-case scenario, colocalization was only agreed by all four methods for 20% of proteins, despite our experiment being constructed to select for cases where colocalization is expected. This suggests caution in performing and interpreting colocalization analyses is warranted.

## INTRODUCTION

Proteins are central to many vital processes^1^, and represent the targets of most drugs. Genome-wide association studies (GWAS) of circulating protein levels have identified many genetic predictors of protein abundance known as protein quantitative trait loci (pQTL)^2^, and integration of pQTL signals with GWAS of diseases has provided critical insights into the molecular pathways underlying diseases^3^. For example, mapping pQTLs onto disease-associated loci has generated *cis*-anchored gene–protein–disease networks that reveal shared biological pathways across diseases and help pinpoint causal genes^4^.

SomaScan and Olink are two high-throughput proteomic platforms widely used for quantifying proteins in biological samples at scale, each leveraging distinct technologies. SomaScan is a Slow Off-rate Modified Aptamer (SOMAmer)-based proteomics assay, which transforms a protein signal into a nucleotide signal by utilizing chemically modified nucleotides. The transformed signal can be detected by relative fluorescence units on microarrays^5^. SomaScan V4.1 measuring up to 7K proteins is used in FinnGen, and 3K SomaScan measuring up to 3K proteins is used in INTERVAL in this investigation.

Olink involves dual antibody binding to the target protein using DNA-labelled antibodies, formation of a unique double-stranded DNA barcode proportional to protein concentration, polymerase chain reaction (PCR) amplification for detection, and readout via quantitative PCR or next-generation sequencing depending on the platform^6^. The Olink Explore 3072 platform considered in this investigation measures more than 2,900 proteins.

Several studies^7–9^ have been published comparing the two platforms, noting only modest correlation between them, and that a considerable proportion of pQTL signals appear to be platform specific. However, good replication of cis-pQTL signals is observed across platforms.

Colocalization is a statistical approach that assesses whether genetic associations for two traits in a specific gene region are driven by the same or distinct causal variants^10^. Enumeration-based colocalization methods, such as coloc^11^ and its multi-signal extension coloc-SuSiE^12^, consider variant-level hypotheses within a Bayesian framework to estimate the posterior probability that two traits share a causal variant. Proportional colocalization approaches, including prop-coloc^13^ and colocPropTest^14^, instead test whether the associations of variants with the two traits are proportional, as implied by colocalization of signals in the region, providing a complementary strategy that potentially is more robust to uncertainty in variant availability.

Enumeration-based approaches have been described as “fine-mapping squared” - they consider the likelihood that each variant in a region is causal for both traits, and compare this to the likelihood that a pair of distinct causal variants are needed to explain the regional associations with the traits, or even that one or both traits have no causal variants in the region. They thus focus on mapping a single causal variant per trait (for coloc), or require a previous step of fine mapping to separate multiple signals in the case of multiple causal variants (for coloc-SuSiE). In both cases, the null hypothesis is no association with either trait. It is possible to test for colocalization across datasets from different ancestries and hence with different linkage disequilibrium (LD) patterns. On the other hand, proportional colocalization methods define the null hypothesis as colocalization, under which, with similar LD in the two datasets, the effects of one or more shared causal variants should be reflected in proportional estimated associations across variants for the two traits. The proportional tests try to be sensitive to departures from this proportional model while maintaining accurate type I error rates under the null. Prop-coloc and colocPropTest differ in that prop-coloc (in its prop-coloc-cond version) tests a single pair of variants, accounting for how those variants are selected, while colocPropTest tests all pairs of variants, accounting for multiple testing. A particular challenge for proportional methods is that true colocalization can be hard to distinguish from lack of power in cases of weak associations. Together, these methods provide several options for investigating shared genetic etiology between traits and diseases and have been widely used. However, the extent to which the methods give reliable evidence in realistic settings is unclear, particularly when, as is common, the two traits are not measured on the same individuals.

In this study, we benchmark these colocalization methods using pQTL data. To provide a baseline scenario, we take a single large dataset, split the dataset in half, and see if protein associations estimated in one half of the dataset colocalize with associations for the same protein estimated in the other half of the dataset. We then consider colocalization using genetic association estimates for the same protein, but measured on different platforms in datasets representing the same population, as well as associations for the same protein measured on the same platform but in datasets representing different populations.

The aim of our investigation is to provide analysts with insight into the practical reliability of various colocalization methods. Colocalization is often used in drug development to triage plausible targets^15^. However, it is unclear how often a successful colocalization finding should be expected when two variants truly share a causal variant. Equally, it is unclear whether factors such as being estimated in datasets representing different populations would lead to failure to colocalize, and so whether a finding indicating non-colocalization (i.e. distinct causal variants) should be taken as strong evidence that the traits do not have a shared etiology in the genetic region considered.

## METHODS

### Overview

We perform colocalization using four different methods on all proteins that have significant associations in the neighborhood of the coding gene region across all four datasets, representing genetic associations with proteins estimated in two UK-based cohorts (one measured using SomaScan, and one using Olink) and a Finnish cohort (proteins measured on both SomaScan and Olink platforms). We investigate colocalization in a range of scenarios, starting with the case where estimates are obtained in two halves of the same dataset, and proceeding to scenarios where the protein associations are obtained in different datasets. In all analyses, we investigate colocalization using genetic association estimates for the same protein (i.e. the two traits are the same protein), but in later scenarios the protein associations are estimated on different platforms or in samples from different populations. For each scenario and each method, we present the proportion of proteins for which a colocalization result was reported, a non-colocalization result was reported, and an unclear result was reported.

More details of our analyses and accompanying software code are available at: https://github.com/sw4rsw4r/pQTL_comparison.git.

### Datasets

The UK Biobank Pharma Proteomics Project (UKB-PPP) is a subset of UK Biobank^16^, a large-scale cohort study of British residents aged 40-69 years at baseline and recruited from 2006-2010. Plasma proteomics were measured on UKB-PPP participants using the Olink Explore 3072 platform. Associations in this study were initially estimated in a discovery cohort comprising 34,557 participants, and were subsequently validated using a replication cohort (n = 17,806). We use associations from the discovery cohort, which consisted exclusively of individuals of European ancestry, making it well-suited for our study objectives. Detailed information on the pQTL analysis performed using these proteomic data is provided in Sun et al (2023)^17^.

The INTERVAL^18,19^ study is a randomized trial of 45,263 British resident blood donors designed to assess whether shortening whole blood donation intervals could optimize blood supply while maintaining donor safety. Plasma proteomics were measured on a subset of INTERVAL participants using the 3K SomaScan assay^20^. Associations used in this study are taken from a genome-wide association analysis of 10.6 million imputed autosomal variants for 2,965 distinct plasma proteins estimated in 3,301 European ancestry participants^21^.

FinnGen is a large-scale public–private initiative that has gathered and analyzed genomic and health information from 500,000 Finnish biobank participants to investigate the genetic underpinnings of human diseases^22^. Proteomics analysis was performed using protein expression profiles measured from two proteomic platforms: SomaScan v4.1 (6,282 proteins measured in 828 individuals) and the Olink Explore 3072 panel (2,897 proteins measured in 619 individuals). The number of proteins shown here is after quality control. The detailed association analysis workflow is described in its webpage^22,23^; in brief, PLINK2 was used to analyze unrelated samples only, and the variant sets differed slightly between the two datasets.

### Colocalization analysis

We assess the performance of four colocalization methods: coloc, coloc-SuSiE, prop-coloc, and colocPropTest. We describe each method in turn.

The coloc method evaluates evidence for five distinct hypotheses regarding the causal variants for two traits in a given genomic region, under the assumption that each trait has no more than one causal variant^11^ in that region. The first hypothesis posits that neither trait has a causal variant in the region (H0). The second and third hypotheses propose that a causal variant exists for only one of the two traits: either the first trait (H1) or the second trait (H2). The fourth hypothesis is that both traits have causal variants, but these variants are distinct and not shared between the traits (H3). The fifth hypothesis is that the two traits share the same causal variant, representing the scenario of colocalization (H4). We interpret a result as “colocalization” if the posterior probability for H4 is greater than 50%, “non-colocalization” if the posterior probability for H3 is greater than 50%, and “insufficient evidence” otherwise.

The coloc-SuSiE (sum of single effects) method is an extension to coloc that first performs fine-mapping using SuSiE to detect credible sets containing causal variants for each trait^24^, and then performs pairwise colocalization using coloc for each pair of credible sets^12^. We interpret a result as “colocalization” if the posterior probability for H4 is greater than 50% for any pair of credible sets, “non-colocalization” if no pair of sets colocalizes and the posterior probability for H3 is greater than 50% for any pair of credible sets, and “insufficient evidence” otherwise.

The prop-coloc method assesses heterogeneity in the error terms of a linear model relating the genetic associations^13^. We here perform prop-coloc-cond, which conditions on the selection of lead variants for each trait to account for bias in variant selection, thereby improving type I error control. The method reports two p-values: one to assess whether the proportionality constant is non-zero (slope test), and the other to assess whether there is heterogeneity. We interpret a result as “colocalization” if p<0.05 for the slope test and p>0.05 for the heterogeneity test, “non-colocalization” if p<0.05 for the slope test and p<0.05 for the heterogeneity test, and “insufficient evidence” otherwise.

The colocPropTest assesses proportionality in trait associations by testing for departure from proportionality of association estimates using all possible variant pairs^25^, focusing on pairs with the smallest marginal p-values when the number of pairs exceeds a threshold (default 10,000). The p-values for each pair of tests are false discovery rate (FDR) corrected, and the minimum FDR corrected p-value is reported, correcting for the number of tests which could have been performed in the case where only a subset of tests are performed. This is equivalent to saying we reject the null hypothesis of proportionality if it is rejected for any pair of variants tested. We interpret a result as “colocalization” if the minimum p>0.05 for the FDR-corrected proportionality test, “non-colocalization” if the minimum p≤0.05 for the FDR-corrected proportionality test, and “insufficient evidence” if the method failed to run (due to insufficient shared variants).

All methods were performed with default settings, including priors of 10^-4^ for *p1* and *p2* and 10^-5^ for *p12* in coloc and coloc-SuSiE. For the prop-coloc method, the presence of highly multicollinear variants can cause the method to fail to run. To prevent such failures, we applied increasingly stringent pruning in each application until a collinearity-induced failure did not occur. Consequently, a single pruning threshold was not applied uniformly across all datasets. As a supplementary analysis, we also present results where the threshold for declaring colocalization and non-colocalization in coloc and coloc-SuSiE is 80% rather than 50%.

### Data preparation

For each protein-coding gene of interest, we extracted pQTL summary statistics within a *cis*-window defined as ±50 kilobases from the transcriptional coordinates (based on Ensembl annotation, GRCh38). Duplicated rsIDs (<0.5% of all variants) due to >2 alleles at the same location were filtered by retaining the record with the greatest association estimate. To enable cross-study comparisons and integrative analyses, we harmonized summary-level pQTL data across datasets. For each target gene region, we considered only variants that were commonly available across all input datasets. Effect and non-effect alleles were aligned across datasets.

The coloc method only requires genetic summary statistics (beta-coefficient and standard error) as inputs, whereas the other three methods require the input of a variant-correlation matrix, also called a linkage disequilibrium (LD) matrix. The coloc-SuSiE method can be run using a common LD matrix, or using separate LD matrices for each trait. LD matrices were obtained from UK Biobank and FinnGen: UK Biobank LD matrix was based on 337K unrelated British-ancestry individuals^26^ and FinnGen LD matrix was based on the Finnish SISu v3 reference panel^27^ (n = 3,775 unrelated Finnish-ancestry individuals), and were harmonized to the same effect and non-effect alleles.

For the baseline scenario (Case 1), we estimated genetic associations ourselves from individual-level data. We divided the UKB-PPP dataset into two equal-sized halves at random, and estimated associations for each variant in the *cis*-gene region (±50 kilobases) using linear regression in PLINK 2.0 for both halves. Covariates were sex, age, and the first 20 principal components of genetic ancestry. Variants with minor allele frequency < 0.001, missing genotype rate > 10%, or Hardy–Weinberg equilibrium p-value < 1×10⁻⁶ were excluded from downstream analyses.

### Protein filtering

In total, 1,140 proteins were commonly present across all four datasets. Proteins with coding regions in the *MHC* region were then removed, leaving 1,132 proteins. Next, proteins mapped to the X chromosome were excluded, resulting in 1,100 proteins. Finally, only proteins having at least one significant signal in the coding gene region with a p-value ≤ 10^-8^ in each platform were retained for downstream analysis, resulting in a final set of 153 proteins used in the analysis (**Figure 1**).

**Figure 1.**
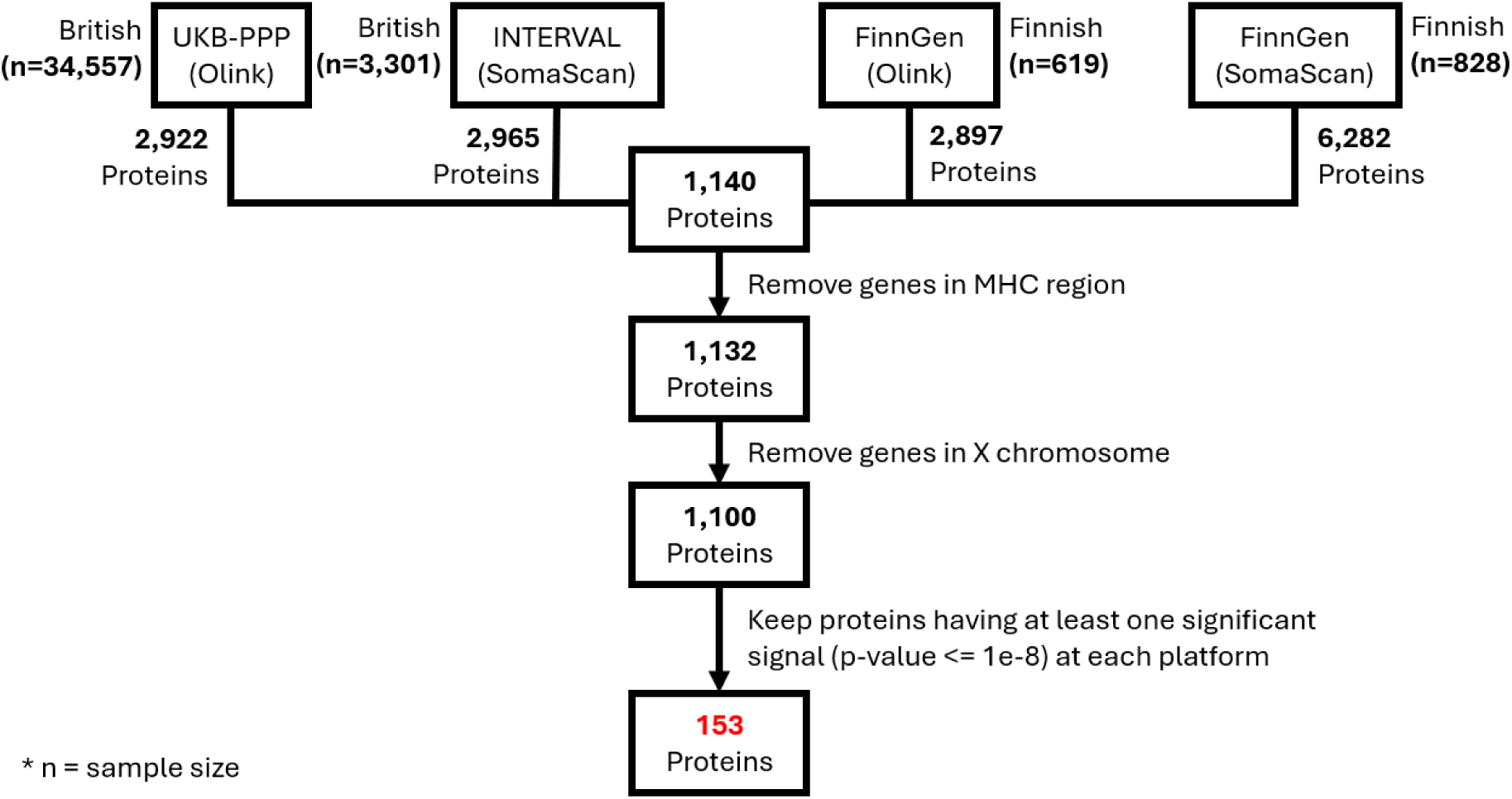
Overview of pQTL datasets and filtering steps used for protein inclusion in the final analysis. For each dataset, the study name, proteomic platform, population, sample size, and the total number of proteins available (post-quality control) are noted. To enable more meaningful comparison, several filtering steps were applied, resulting in a final set of 153 proteins used for analysis.

### Comparison scenarios

The key information for the scenarios in the comparison analysis is described in **Table 1**, with detailed information for each case provided below.

- Case 1 (Baseline scenario: same dataset): genetic associations were taken from separate halves of UKB-PPP (Olink), and the LD matrix was taken from UK Biobank.
- Case 2 (Same population, different platforms):

- Case 2B: genetic associations were taken from UK Biobank (Olink) and INTERVAL (SomaScan), and the LD matrix was taken from UK Biobank.
- Case 2F: genetic associations were taken from FinnGen (Olink and SomaScan), and the LD matrix was taken from FinnGen.
- Case 3 (Different populations, Olink platform):

- Case 3O: genetic associations were taken from UKB-PPP (Olink) and FinnGen (Olink). The LD matrix was taken from UK Biobank (Case 3OB), FinnGen (Case 3OF), or matched to each dataset (Case 3OM; note this is only possible for coloc-SuSiE, as other methods only allow a single LD matrix).
- Case 4 (Different populations, SomaScan platform):

- Case 4S: genetic associations were taken from INTERVAL (SomaScan) and FinnGen (SomaScan). The LD matrix was taken from UK Biobank (Case 4SB), FinnGen (Case 4SF), or matched to each dataset (Case 4SM).

**Table 1.** Summary of scenarios in comparative analysis.

|  | Population | Study | Platform | LD matrix |  | Population | Study | Platform | LD matrix |
| --- | --- | --- | --- | --- | --- | --- | --- | --- | --- |
| **CASE 1** | British | UKB-PPP | OLINK | UKB | **vs** | British | UKB-PPP | OLINK | UKB |
| **CASE 2-B** | British | UKB-PPP | OLINK | UKB |  | British | INTERVAL | SomaScan | UKB |
| **CASE 2-F** | Finnish | FinnGen | OLINK | FinnGen |  | Finnish | FinnGen | SomaScan | FinnGen |
| **CASE 3-OM** | British | UKB-PPP | OLINK | UKB |  | Finnish | FinnGen | OLINK | FinnGen |
| **CASE 3-OF** | British | UKB-PPP | OLINK | FinnGen |  | Finnish | FinnGen | OLINK | FinnGen |
| **CASE 3-OB** | British | UKB-PPP | OLINK | UKB |  | Finnish | FinnGen | OLINK | UKB |
| **CASE 4-SM** | British | INTERVAL | SomaScan | UKB |  | Finnish | FinnGen | SomaScan | FinnGen |
| **CASE 4-SF** | British | INTERVAL | SomaScan | FinnGen |  | Finnish | FinnGen | SomaScan | FinnGen |
| **CASE 4-SB** | British | INTERVAL | SomaScan | UKB |  | Finnish | FinnGen | SomaScan | UKB |

As coloc does not require an LD matrix, only one version of the coloc method was run in Cases 3 and 4. Two versions of prop-coloc and colocPropTest were run for each case (British and Finnish LD matrices), and three versions of coloc-SuSiE (British, Finnish, and matched LD matrices).

## RESULTS

Results are summarized in **Table 2** (summary of results for each method and scenario) and **Figure 2** (UpSet plot showing agreement between methods) where in each case, only the top five patterns are presented. More detailed information is provided in **Supplementary Tables 1-9**, which show the pairwise agreement between methods in each scenario. Similar findings are shown in **Supplementary Table 10**, which displays results from coloc and coloc-SuSiE with an 80% threshold for declaring colocalization and non-colocalization. In coloc-SuSiE, more than one credible set was found for over 95% of proteins across each dataset.

**Figure 2.**
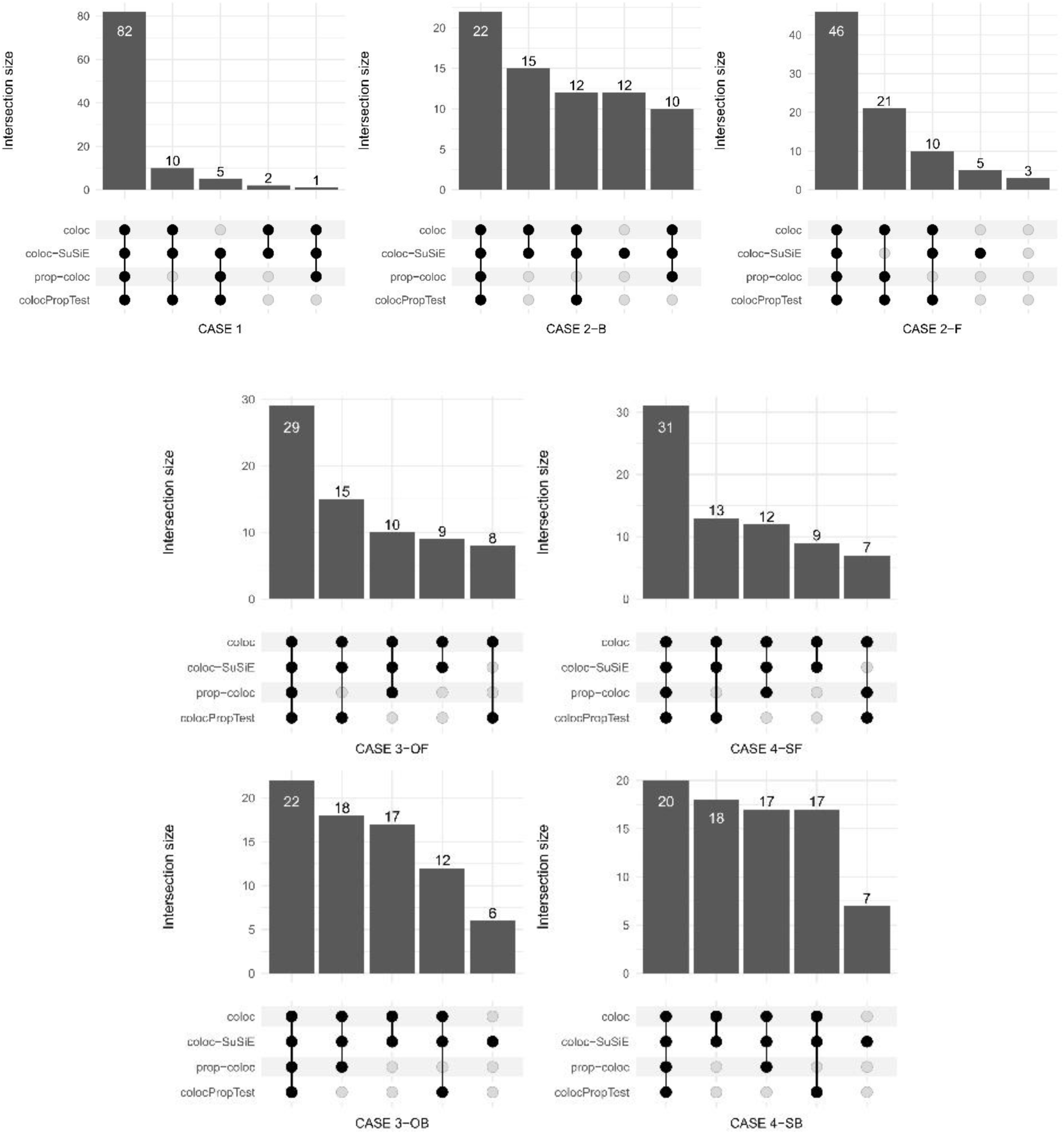
UpSet plot visualizations showing patterns of colocalization results across four methods in each scenario. Only the top 5 patterns are shown in each case. Numbers represent the proportion of proteins fitting each pattern (%).

**Table 2.** Summary of findings across methods and scenarios: proportion of proteins with evidence for colocalization and non-colocalization, and with insufficient evidence for either conclusion.

|  | coloc |  |  | coloc-SuSiE |  |  | prop-coloc |  |  | colocPropTest |  |  |
| --- | --- | --- | --- | --- | --- | --- | --- | --- | --- | --- | --- | --- |
|  | C | NC | IS | C | NC | IS | C | NC | IS | C | NC | IS |
| CASE 1 | 0.95 | 0.05 | 0.00 | 1.00 | 0.00 | 0.00 | 0.88 | 0.12 | 0.00 | 0.97 | 0.03 | 0.00 |
| CASE 2-B | 0.67 | 0.33 | 0.00 | 0.84 | 0.09 | 0.07 | 0.50 | 0.40 | 0.10 | 0.45 | 0.55 | 0.00 |
| CASE 2-F | 0.80 | 0.20 | 0.00 | 0.71 | 0.03 | 0.25 | 0.76 | 0.22 | 0.03 | 0.86 | 0.14 | 0.00 |
| CASE 3-OM | 0.83 | 0.16 | 0.01 | 0.58 | 0.15 | 0.27 |  |  |  |  |  |  |
| CASE 3-OF |  |  |  | 0.72 | 0.01 | 0.27 | 0.56 | 0.43 | 0.01 | 0.65 | 0.35 | 0.00 |
| CASE 3-OB |  |  |  | 0.81 | 0.09 | 0.10 | 0.56 | 0.33 | 0.11 | 0.43 | 0.57 | 0.00 |
| CASE 4-SM | 0.79 | 0.20 | 0.01 | 0.65 | 0.19 | 0.16 |  |  |  |  |  |  |
| CASE 4-SF |  |  |  | 0.81 | 0.01 | 0.18 | 0.65 | 0.30 | 0.05 | 0.61 | 0.38 | 0.01 |
| CASE 4-SB |  |  |  | 0.90 | 0.08 | 0.03 | 0.50 | 0.39 | 0.11 | 0.48 | 0.52 | 0.01 |

\*C: colocalization, NC: non-colocalization, IS: Insufficient evidence,

### Case 1: Baseline scenario, same dataset

In Case 1, we expect colocalization in all analyses. This was achieved by coloc-SuSiE, which had a 100% success rate. Amongst other methods, colocPropTest declared colocalization for 97%, coloc for 95%, and prop-coloc for 88% of proteins. For all proteins, at least two methods agreed there was colocalization.

### Case 2: Same population, different platform

In Cases 2 to 4, we hope to see high levels of colocalization, but results may not achieve 100% for many reasons, including platform-specific artefacts and differences in genetic architecture between populations. However, such differences are likely to be common in practice, and so we want to know if the methods are robust to these deviations.

In Case 2B (British), coloc-SuSiE declares colocalization for 84% of proteins and non-colocalization for 9% of proteins. Other methods are less confident, with colocalization declared for 45-67% of proteins, and non-colocalization declared for 33-55% of proteins. All four methods only agree on colocalization for 22% of proteins.

In Case 2F (Finnish), coloc-SuSiE declares colocalization for 71% of proteins and non-colocalization for 3% of proteins. Other methods are similarly confident, with colocalization declared for 76-86% of proteins, and non-colocalization declared for 14-22% of proteins. All four methods agree on colocalization for 46% of proteins.

### Cases 3 and 4: Same platform, different population

In Cases 3 and 4, coloc declares colocalization for 83% and 79% of proteins, and non-colocalization for 16% and 20% of proteins. With matched LD matrices, coloc-SuSiE does less well, declaring colocalization for 58% and 65% of proteins, and non-colocalization for 15% and 19% of proteins. Oddly, coloc-SuSiE performs much better with a single LD matrix, whether this is taken from UK Biobank (81% or 90% colocalization) or FinnGen (72% or 81% colocalization). The prop-coloc and colocPropTest methods perform similarly in these scenarios, declaring colocalization for 43-65% of proteins, and non-colocalization for 30-57% of proteins. All four methods agree on colocalization for 20-31% of proteins.

We note that proportional colocalization methods require the datasets to have the same LD structure. We applied these methods here to assess their performance in practice with datasets with somewhat different LD structures (both datasets comprise European ancestry individuals, but the genetic architecture of Finnish Europeans is distinct from other European ancestry populations).

## Discussion

In our comparative colocalization analysis, we observed the proportion of findings indicating colocalization dropping well below 100% in all but the baseline scenario, and with substantial numbers of findings indicating non-colocalization. Although some of these non-colocalization findings may be explained by genuine differences in genetic architecture between UK and Finnish populations, some may reflect true biological or technical differences between the measurement technologies employed. We also found a high degree of disagreement between colocalization methods. Enumeration methods tended to outperform proportional methods in most scenarios. However, performance was context-dependent, and no single approach dominated across all scenarios, with coloc-SuSiE reporting the highest rate of colocalization in Case 1, Case 2B, and Case 4; colocPropTest in Case 2F; and coloc in Case 3.

There are many ways in which this analysis does not treat the various methods fairly. The enumeration colocalization methods (coloc and coloc-SuSiE) operate in a Bayesian framework, and so only declare colocalization if there is sufficient evidence to support colocalization. In contrast, for the proportional colocalization methods, the null hypothesis is colocalization, and so the methods declare colocalization in the absence of evidence against colocalization. This may explain the better performance of proportional colocalization methods in Case 2F compared to Case 2B. Case 2F uses the correct Finnish LD for both datasets, while case 2B uses UK Biobank LD for the INTERVAL cohort. Small differences in LD between the datasets can lead to deviation from proportionality, and hence rejection of the colocalisation null hypothesis, particularly given the substantially larger sample sizes (leading to increased power to reject colocalization) available in case 2B. Additionally, coloc-SuSiE declares colocalization if at least 1 pair of credible sets indicates colocalization, meaning that coloc-SuSiE is more likely to declare colocalization in ambiguous cases and cases with multiple causal variants. Finally, we say that a method performed “better” if it declares colocalization more often, as the ground truth in all scenarios considered is colocalization. In all scenarios other than Case 1, there are genuine reasons why estimates may not colocalize. If a method is biased towards declaring colocalization, it may appear to perform better in this study than a method that is correctly detecting deviation from colocalization.

One strange result is that coloc-SuSiE appears to perform better when the LD matrices are misspecified rather than correctly specified. This may be because using the same LD matrix for both traits increases the likelihood of identifying the same causal variants, and hence declaring colocalization. Alternatively, there may be some other reason for a bias towards colocalization in this setting. Hence, the method may not be performing better, but it appears to be performing better because of a skew towards reporting colocalization.

Our aim in this investigation was to provide a comparison of methods as they are likely to be implemented in practice, and we have prioritized this over taking decisions that may be taken in practice to ensure “fairness” between methods. We therefore discourage readers from drawing strong conclusions about which method performs best, and instead to focus on the evidence provided by each method, and the extent to which this should encourage (or discourage) a conclusion of shared aetiology between the traits. We found that platform, population, method, and choice of LD matrix can markedly affect colocalization outcomes, with concordance between all four methods supporting colocalization reaching as low as 20% in the most discordant scenario. Therefore, colocalization analyses should be applied and interpreted with caution, particularly when the datasets used for the different traits differ considerably. Particular caution should be exercised when attempting to colocalize datasets from different ancestries. While coloc can be robust to ancestry differences, LD mismatch will typically lead to departure from proportionality in the proportional colocalization methods, and as suggested above using an incorrect LD matrix may skew the results of coloc-SuSiE.

From an optimistic perspective, all methods performed well in Case 1, with coloc-SuSiE declaring colocalization for 100% of proteins, and at most one method disagreeing in the majority of examples. This shows the value of triangulation of evidence from multiple methods; if different methods disagree, this provides caution that the verdict of colocalization or non-colocalization may be uncertain. Additionally, the coloc and coloc-SuSiE methods generally performed well, with coloc-SuSiE often declaring “insufficient evidence” rather than non-colocalization. However, from a pessimistic perspective, aside from in Case 1, all methods declared non-colocalization for around 20% of proteins in at least one scenario. This should calibrate our intuition that when we see a non-colocalization finding in practice, we should retain some skepticism in whether this finding can be fully trusted.

There are several limitations to this work. As stated above, there is no way to make this a fair comparison^28^, and our design favors methods that more strongly tend to declare colocalization. There is no obvious way to construct negative scenarios other than by simulating synthetic data, whereas we want to focus on real-life examples. We only considered datasets with protein data. On the one hand, this may favor the detection of colocalization, as protein associations are typically strong. However, given the complexity of proteins, this may make it more difficult to detect colocalization, as many proteins may have multiple hits in their coding gene regions. If different variants affect different aspects of protein biology in a way that interacts with assay binding, we may see departures from proportionality in the genetic associations. The proportional colocalization methods also assume the same genetic architecture in associations with both traits, which is unlikely to hold when they come from different populations. However, we deliberately wanted to investigate the performance of methods under these conditions, as they occur frequently in practice. Our results suggest that the proportional colocalization methods do not perform well when the assumption of equal LD in the two datasets is violated. A further limitation is that we only performed analyses on datasets comprising European ancestry individuals, and we did not consider scenarios where the datasets represent different ancestral populations, which is likely to lead to greater discrepancies than those between British and Finnish populations presented here.

In conclusion, while all methods were well-calibrated in the baseline scenario, they struggled to different degrees when the datasets varied in terms of platform and population. Colocalization can be a valuable tool for triaging and prioritizing trait pairs as having evidence for shared etiology. However, like all statistical methods, it has limitations, and results should not be thought of as unquestionable truth.

## Acknowledgements

S. B. is supported by the Wellcome Trust (225790/Z/22/Z) and the United Kingdom Research and Innovation Medical Research Council (MRC) (MC_UU_00040/01). C. W. is supported by the Wellcome Trust (WT 220788) and MRC (MC_UU_00040/01). J. P. is supported by a Gates Cambridge Scholarship.

## Code availability

R scripts to perform all case studies in this manuscript including data acquisition, data preprocessing, colocalization analysis are available at https://github.com/sw4rsw4r/pQTL_comparison and R Markdown document to enable readers to easily reproduce and verify the analysis is available at https://sw4rsw4r.github.io/pQTL_comparison/pQTL_comparison.html

## Ethics declarations

Most analyses in our study were conducted using publicly available summarized data, for which specific ethical approval was not required. Ethical approval details for the original studies can be found in the relevant references. In addition, for a subset of analyses that utilized individual-level genotype data from the UK Biobank, we generated and analyzed only summary-level data that contained no personally identifiable information. Therefore, these analyses also do not require separate ethical approval.

---

**Supplementary Table 1.** Method-by-method concordance of colocalization classifications (CASE 1)

| CASE 1 |  | coloc-SuSiE |  |  | prop-coloc |  |  | colocPropTest |  |  |
| --- | --- | --- | --- | --- | --- | --- | --- | --- | --- | --- |
|  |  | C | NC | IS | C | NC | IS | C | NC | IS |
| coloc | C | 0.95 | 0.00 | 0.00 | 0.83 | 0.12 | 0.00 | 0.92 | 0.03 | 0.00 |
|  | NC | 0.05 | 0.00 | 0.00 | 0.05 | 0.00 | 0.00 | 0.05 | 0.00 | 0.00 |
|  | IS | 0.00 | 0.00 | 0.00 | 0.00 | 0.00 | 0.00 | 0.00 | 0.00 | 0.00 |
| coloc-SuSiE | C |  |  |  | 0.88 | 0.12 | 0.00 | 0.97 | 0.03 | 0.00 |
|  | NC |  |  |  | 0.00 | 0.00 | 0.00 | 0.00 | 0.00 | 0.00 |
|  | IS |  |  |  | 0.00 | 0.00 | 0.00 | 0.00 | 0.00 | 0.00 |
| prop-coloc | C |  |  |  |  |  |  | 0.86 | 0.01 | 0.00 |
|  | NC |  |  |  |  |  |  | 0.10 | 0.02 | 0.00 |
|  | IS |  |  |  |  |  |  | 0.00 | 0.00 | 0.00 |

**Supplementary Table 2.** Method-by-method concordance of colocalization classifications (CASE 2-B)

| CASE 2-B |  | coloc-SuSiE |  |  | prop-coloc |  |  | colocPropTest |  |  |
| --- | --- | --- | --- | --- | --- | --- | --- | --- | --- | --- |
|  |  | C | NC | IS | C | NC | IS | C | NC | IS |
| coloc | C | 0.59 | 0.03 | 0.05 | 0.35 | 0.25 | 0.07 | 0.39 | 0.28 | 0.00 |
|  | NC | 0.24 | 0.07 | 0.03 | 0.16 | 0.14 | 0.03 | 0.07 | 0.27 | 0.00 |
|  | IS | 0.00 | 0.00 | 0.00 | 0.00 | 0.00 | 0.00 | 0.00 | 0.00 | 0.00 |
| coloc-SuSiE | C |  |  |  | 0.42 | 0.33 | 0.08 | 0.37 | 0.47 | 0.00 |
|  | NC |  |  |  | 0.05 | 0.04 | 0.01 | 0.04 | 0.05 | 0.00 |
|  | IS |  |  |  | 0.03 | 0.03 | 0.01 | 0.05 | 0.03 | 0.00 |
| prop-coloc | C |  |  |  |  |  |  | 0.29 | 0.22 | 0.00 |
|  | NC |  |  |  |  |  |  | 0.14 | 0.26 | 0.00 |
|  | IS |  |  |  |  |  |  | 0.03 | 0.07 | 0.00 |

**Supplementary Table 3.** Method-by-method concordance of colocalization classifications (CASE 2-F)

| CASE 2-F |  | coloc-SuSiE |  |  | prop-coloc |  |  | colocPropTest |  |  |
| --- | --- | --- | --- | --- | --- | --- | --- | --- | --- | --- |
|  |  | C | NC | IS | C | NC | IS | C | NC | IS |
| coloc | C | 0.58 | 0.01 | 0.21 | 0.67 | 0.12 | 0.01 | 0.78 | 0.03 | 0.00 |
|  | NC | 0.13 | 0.02 | 0.05 | 0.08 | 0.10 | 0.01 | 0.08 | 0.11 | 0.00 |
|  | IS | 0.00 | 0.00 | 0.00 | 0.00 | 0.00 | 0.00 | 0.00 | 0.00 | 0.00 |
| coloc-SuSiE | C |  |  |  | 0.52 | 0.18 | 0.02 | 0.63 | 0.08 | 0.00 |
|  | NC |  |  |  | 0.03 | 0.00 | 0.01 | 0.02 | 0.01 | 0.00 |
|  | IS |  |  |  | 0.22 | 0.04 | 0.00 | 0.22 | 0.04 | 0.00 |
| prop-coloc | C |  |  |  |  |  |  | 0.71 | 0.05 | 0.00 |
|  | NC |  |  |  |  |  |  | 0.14 | 0.08 | 0.00 |
|  | IS |  |  |  |  |  |  | 0.01 | 0.01 | 0.00 |

**Supplementary Table 4.** Method-by-method concordance of colocalization classifications (CASE 3-OM)

| CASE 3-OM |  | coloc-SuSiE |  |  |
| --- | --- | --- | --- | --- |
|  |  | C | NC | IS |
| coloc | C | 0.49 | 0.14 | 0.20 |
|  | NC | 0.08 | 0.01 | 0.07 |
|  | IS | 0.00 | 0.00 | 0.01 |

**Supplementary Table 5.** Method-by-method concordance of colocalization classifications (CASE 3-OF)

| CASE 3-OF |  | prop-coloc |  |  | colocPropTest |  |  |
| --- | --- | --- | --- | --- | --- | --- | --- |
|  |  | C | NC | IS | C | NC | IS |
| coloc-SuSiE | C | 0.42 | 0.29 | 0.01 | 0.46 | 0.26 | 0.00 |
|  | NC | 0.01 | 0.01 | 0.00 | 0.01 | 0.00 | 0.00 |
|  | IS | 0.13 | 0.13 | 0.01 | 0.18 | 0.09 | 0.00 |
| prop-coloc | C |  |  |  | 0.38 | 0.18 | 0.00 |
|  | NC |  |  |  | 0.25 | 0.18 | 0.00 |
|  | IS |  |  |  | 0.01 | 0.00 | 0.00 |

**Supplementary Table 6.** Method-by-method concordance of colocalization classifications (CASE 3-OB)

| CASE 3-OB |  | prop-coloc |  |  | colocPropTest |  |  |
| --- | --- | --- | --- | --- | --- | --- | --- |
|  |  | C | NC | IS | C | NC | IS |
| coloc-SuSiE | C | 0.46 | 0.28 | 0.07 | 0.35 | 0.46 | 0.00 |
|  | NC | 0.06 | 0.01 | 0.02 | 0.03 | 0.06 | 0.00 |
|  | IS | 0.04 | 0.04 | 0.02 | 0.05 | 0.05 | 0.00 |
| prop-coloc | C |  |  |  | 0.27 | 0.28 | 0.00 |
|  | NC |  |  |  | 0.10 | 0.24 | 0.00 |
|  | IS |  |  |  | 0.06 | 0.05 | 0.00 |

**Supplementary Table 7.** Method-by-method concordance of colocalization classifications (CASE 4-SM)

| CASE 4-SM |  | coloc-SuSiE |  |  |
| --- | --- | --- | --- | --- |
|  |  | C | NC | IS |
| coloc | C | 0.50 | 0.16 | 0.13 |
|  | NC | 0.15 | 0.03 | 0.02 |
|  | IS | 0.00 | 0.00 | 0.01 |

**Supplementary Table 8.** Method-by-method concordance of colocalization classifications (CASE 4-SF)

| CASE 4-SF |  | prop-coloc |  |  | colocPropTest |  |  |
| --- | --- | --- | --- | --- | --- | --- | --- |
|  |  | C | NC | IS | C | NC | IS |
| coloc-SuSiE | C | 0.54 | 0.24 | 0.03 | 0.50 | 0.30 | 0.01 |
|  | NC | 0.01 | 0.00 | 0.01 | 0.00 | 0.01 | 0.00 |
|  | IS | 0.11 | 0.06 | 0.01 | 0.11 | 0.07 | 0.00 |
| prop-coloc | C |  |  |  | 0.44 | 0.21 | 0.01 |
|  | NC |  |  |  | 0.16 | 0.14 | 0.00 |
|  | IS |  |  |  | 0.01 | 0.03 | 0.00 |

**Supplementary Table 9.** Method-by-method concordance of colocalization classifications (CASE 4-SB)

| CASE 4-SB |  | prop-coloc |  |  | colocPropTest |  |  |
| --- | --- | --- | --- | --- | --- | --- | --- |
|  |  | C | NC | IS | C | NC | IS |
| coloc-SuSiE | C | 0.46 | 0.35 | 0.09 | 0.44 | 0.45 | 0.01 |
|  | NC | 0.04 | 0.03 | 0.01 | 0.03 | 0.05 | 0.00 |
|  | IS | 0.01 | 0.01 | 0.01 | 0.01 | 0.02 | 0.00 |
| prop-coloc | C |  |  |  | 0.27 | 0.22 | 0.01 |
|  | NC |  |  |  | 0.14 | 0.24 | 0.00 |
|  | IS |  |  |  | 0.06 | 0.05 | 0.00 |

**Supplementary Table 10.**
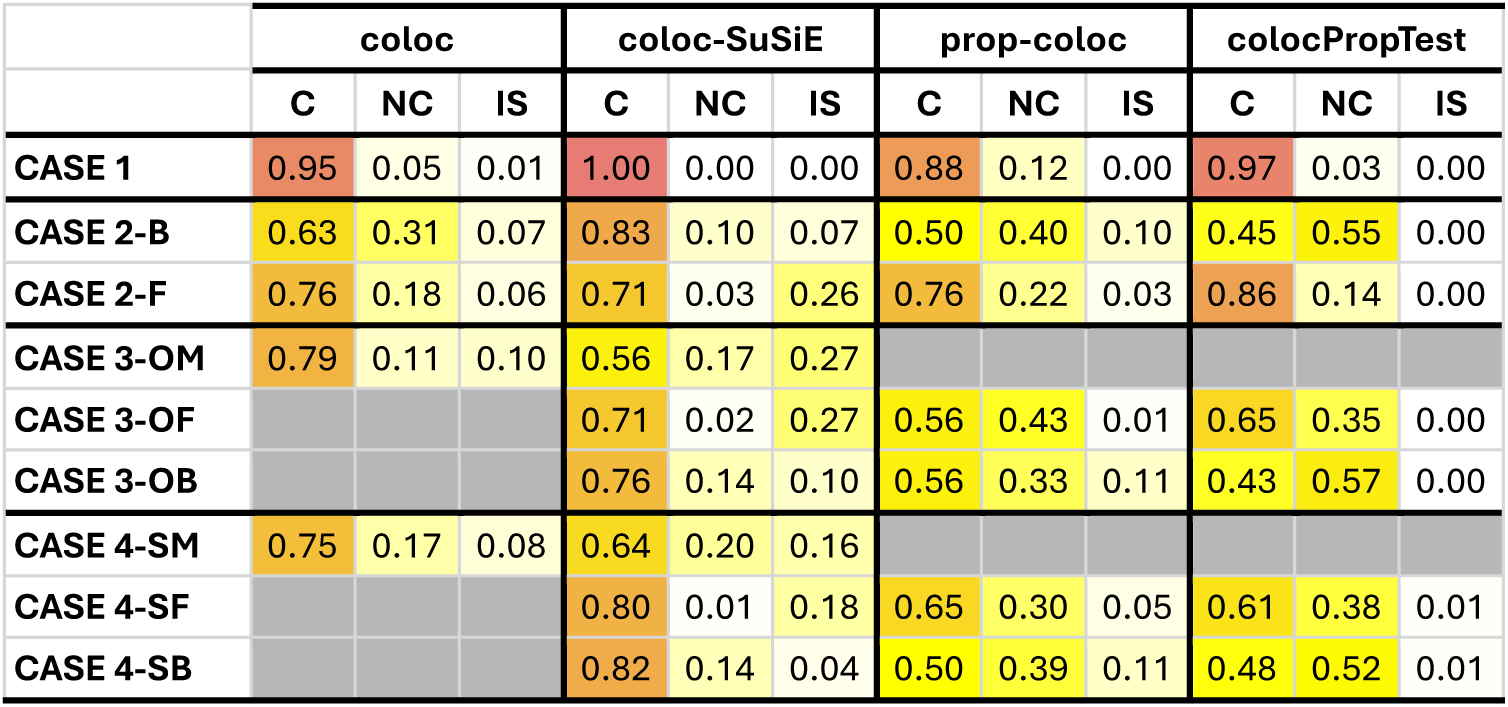
Percentages of colocalization outcomes across methods and scenarios with different posterior probability threshold (0.8) for coloc and coloc-SuSiE. All other criteria were identical to those described in Table 2, except that the posterior probability thresholds applied to H3 and H4 in coloc and coloc-SuSiE were set to 0.8.

## References

1. Liang, J., Tian, J., Zhang, H., Li, H. & Chen, L. Proteomics: An In-Depth Review on Recent Technical Advances and Their Applications in Biomedicine. Med Res Rev (2025).

2. Deng, Y.-T. et al. Atlas of the plasma proteome in health and disease in 53,026 adults. Cell 188, 253–271 (2025).

3. Ferkingstad, E. et al. Large-scale integration of the plasma proteome with genetics and disease. Nat Genet 53, 1712–1721 (2021).

4. Pietzner, M. et al. Mapping the proteo-genomic convergence of human diseases. Science (1979) 374, eabj1541 (2021).

5. Sattlecker, M. et al. Alzheimer’s disease biomarker discovery using SOMAscan multiplexed protein technology. Alzheimer’s & Dementia 10, 724–734 (2014).

6. Wik, L. et al. Proximity extension assay in combination with next-generation sequencing for high-throughput proteome-wide analysis. Molecular & Cellular Proteomics 20, 100168 (2021).

7. Rooney, M. R., et al. Plasma proteomic comparisons change as coverage expands for SomaLogic and Olink. Medrxiv (2024).

8. Katz, D. H. et al. Proteomic profiling platforms head to head: leveraging genetics and clinical traits to compare aptamer-and antibody-based methods. Sci Adv 8, eabm5164 (2022).

9. Eldjarn, G. H. et al. Author Correction: Large-scale plasma proteomics comparisons through genetics and disease associations. Nature 630, E3–E3 (2024).

10. Zuber, V. et al. Combining evidence from Mendelian randomization and colocalization: Review and comparison of approaches. The American Journal of Human Genetics 109, 767–782 (2022).

11. Giambartolomei, C. et al. Bayesian test for colocalisation between pairs of genetic association studies using summary statistics. PLoS Genet 10, e1004383 (2014).

12. Wallace, C. A more accurate method for colocalisation analysis allowing for multiple causal variants. PLoS Genet 17, e1009440 (2021).

13. Patel, A., Whittaker, J. C. & Burgess, S. A frequentist test of proportional colocalization after selecting relevant genetic variants. arXiv preprint arXiv:2402.12171 (2024).

14. Wallace, C., Robins, C. & Johnson, T. Variable information across SNPs in GWAS data can cause false rejections of colocalisation which can be resolved by proportional colocalisation tests. bioRxiv 2025–2029 (2025).

15. Burgess, S. et al. Using genetic association data to guide drug discovery and development: Review of methods and applications. The American Journal of Human Genetics 110, 195–214 (2023).

16. Sudlow, C. et al. UK biobank: an open access resource for identifying the causes of a wide range of complex diseases of middle and old age. PLoS Med 12, e1001779 (2015).

17. Sun, B. B. et al. Plasma proteomic associations with genetics and health in the UK Biobank. Nature 622, 329–338 (2023).

18. Di Angelantonio, E. et al. Efficiency and safety of varying the frequency of whole blood donation (INTERVAL): a randomised trial of 45 000 donors. The Lancet 390, 2360–2371 (2017).

19. Moore, C. et al. The INTERVAL trial to determine whether intervals between blood donations can be safely and acceptably decreased to optimise blood supply: study protocol for a randomised controlled trial. Trials 15, 363 (2014).

20. Tanaka, T. et al. Plasma proteomic biomarker signature of age predicts health and life span. Elife 9, e61073 (2020).

21. Sun, B. B. et al. Genomic atlas of the human plasma proteome. Nature 558, 73–79 (2018).

22. Kurki, M. I. et al. FinnGen provides genetic insights from a well-phenotyped isolated population. Nature 613, 508–518 (2023).

23. FinnGen. FinnGen Documentation.

24. Zou, Y., Carbonetto, P., Wang, G. & Stephens, M. Fine-mapping from summary data with the “Sum of Single Effects” model. PLoS Genet 18, e1010299 (2022).

25. Chris Wallace. colocPropTest. Preprint at https://cran.r-project.org/web/packages/colocPropTest/index.html (2024).

26. Weissbrod, O. et al. Functionally informed fine-mapping and polygenic localization of complex trait heritability. Nat Genet 52, 1355–1363 (2020).

27. Kanai, M. et al. Insights from complex trait fine-mapping across diverse populations. medrxiv 2021–2029 (2021).

28. Boulesteix, A.-L., Lauer, S. & Eugster, M. J. A. A plea for neutral comparison studies in computational sciences. PLoS One 8, e61562 (2013).

